# Neuroprotective Parkinson’s Disease Therapeutic: Transition Metal Dichalcogenide Nanoflower Treatments Alleviate Pathological Cell Stress

**DOI:** 10.1101/2025.09.29.679305

**Authors:** Charles L. Mitchell, Mikhail Matveyenka, Harris C. Brown, Jessica Aldape, Payton Moore, Kha-Tran Nguyen, John C. Walker, Bryce Pearson, Joshua Skrehot, Dmitry Kurouski

## Abstract

Parkinson’s disease (PD) is triggered by irreversible degeneration of dopaminergic neurons in the midbrain, hypothalamus, and thalamus. Although the underlying molecular etiology of these pathological processes remains unclear, progressive aggregation of alpha-synuclein (α-syn) and mitochondrial dysfunction are two expected mechanisms implicated in neuronal degeneration. Accumulating evidence indicates that transition metal dichalcogenide (TMD) nanoflowers (NFs), a novel class of nanomaterials, can restore mitochondrial health by the activation of mitochondrial biogenesis. However, therapeutic potential of TMD NFs in PD remains unclear. The current study investigates the neuroprotective properties of molybdenum disulfide (MoS_2_) and molybdenum diselenide (MoSe_2_) nanoflowers (NFs) in neurons and astrocytes exposed to α-syn aggregates. It was found that MoS_2_ and MoSe_2_ suppressed α-syn-induced unfolded protein response (UPR) in the endoplasmic reticulum, and upregulated autophagy and exocytosis of α-syn fibrils. TMD NFs also reversed α-syn-induced damage of cell mitochondria, simultaneously stimulating mitochondrial biogenesis. As a result, a drastic decrease in ROS levels in both neurons and astrocytes was observed. These results show that MoS_2_ or MoSe_2_ NFs could fully rescue neurons and astrocytes from the cytotoxic effects of α-syn fibrils. Neuroprotective properties of these novel nanomaterials were further explored in *Caenorhabditis elegans* that overexpress α-syn. Nematodes that received NFs experienced a drastic reduction in the amount of aggregated α-syn which resulted in a significant increase in *C. elegans* lifespan. These findings indicated that MoS_2_ or MoSe_2_ NFs could be used as novel therapeutic to decelerate the progression of PD.

## INTRODUCTION

Parkinson’s disease (PD) is the fastest increasing neurological disorder, with an estimated count of 25.2 million diagnoses worldwide by 2050^1^. In the United States alone, there are approximately 60,000 cases of PD diagnosed annually, accounting for a share of the medical market worth at least $52 billion^2^. PD is characterized by the progressive degeneration of dopaminergic neurons in the midbrain, hypothalamus, and thalamus.^3^ This gives rise to numerous PD symptoms, including bradykinesia, postural and rest tremors, rigidity of the muscles, and postural instability^4, 5^, albeit symptoms are heterogeneous between patients^6^. The etiology of PD has yet to be completely understood, however, a growing body of evidence suggests that the onset and propagation of neurodegeneration are triggered by the abrupt aggregation of alpha synuclein (α-syn) and progressive mitochondrial dysfunction ^7-9^.

α-Syn is a low molecular weight protein consisting of 140 amino acid residues that is encoded by the chromosome 4 *SNCA* gene^10^. Although the exact physiological function of α-syn remains unclear, it is known that α-syn facilitates trafficking of presynaptic vesicles releasing neurotransmitters into the synaptic cleft^11, 12^,^13, 14^. α-Syn is primarily localized in the membranes of presynaptic terminals of dopaminergic neurons^15^, although it is also present in many organelles, such as the mitochondria^16^, the endoplasmic reticulum^17^, Golgi apparatus^18, 19^, and nuclei^20^. Although cytosolic α-syn is an intrinsically disordered protein, it folds into an α-helical structure upon embedding into the lipid membranes^21-23^. Under pathological conditions, α-syn aggregates into β-sheet-rich oligomers and fibrils^24, 25^. This conformational transition of α-syn is implicated with the onset and progression of PD^26, 27^. Along with the accumulation of toxic protein aggregates, PD pathology heavily involves mitochondrial dysfunction^28-31^. Under pathological conditions, mitochondria produce an excessive amount of reactive oxygen species (ROS), experience a detrimental depolarization of membrane potential, and a decrease in the production of ATP^32^. Mitochondrial dysfunction further exacerbates cell malfunction, which results in the degeneration of both neurons and astrocytes as observed in PD.

Decades of research have investigated the efficacy of therapeutic approaches to treat PD, primarily focused on inhibiting amyloid fibril formation, reducing amyloid burden in the brain, and alleviating mitochondrial dysfunction. For instance, numerous attempts have been made to inhibit α-syn aggregation using small molecular candidates^33^, natural polyvinyls^34^, polyphenols^35-37^, antioxidants^38, 39^ and protein chaperones^40^. Several research groups explored the possibility of utilizing antibodies, such as Cinpanemab^41^ and Prasinezumab^42^, to reduce the burden of cytotoxic α-syn aggregates in the brain^43^. Finally, therapeutic properties of Coenzyme Q_10_ ^38^ and Idebenone^44, 45^ in alleviation of mitochondrial dysfunction were explored. However, these strategies did not reach expected neuroprotection allowing only to alleviate PD symptoms. These limitations catalyzed the search for novel neuroprotective therapeutic approaches that can be used to decelerate the progression of PD.

Transition metal dichalcogenides (TMD) are a class of nanomaterials with unique optical, catalytic, and biological properties.^46-48^ TMDs are comprised of a transition metal atom, such as molybdenum (Mo) or tungsten (W), coordinated with two chalcogenide atoms of sulfur (S) or selenium (Se).^49^ Hydrothermal synthesis enables self-assembly of metal and chalcogenide atoms into nanoflakes that subsequently organize into nanoflowers (NFs), supramolecular aggregates with high surface area to volume ratio. TMDs, including molybdenum disulfide (MoS_2_) NFs, exhibit unique catalase, superoxide dismutase, and peroxidase properties^50^. Furthermore, it has been shown that both MoS_2_ and molybdenum diselenide (MoSe_2_) NFs facilitated mitochondrial biogenesis and reduced levels of ROS in neurons and astrocytes.^51^ These and other biological properties of TMD NFs open new avenues for the utilization of these nanomaterials as therapies for cancer,^52^ inflammation,^53^ and Alzheimer’s disease (AD)^54^ treatment. However, therapeutic potential of TMD NFs in PD remains unclear.

This study investigates neuroprotective properties of MoS_2_ and MoSe_2_ NFs. The results show that both TMD NFs fully rescued neurons and astrocytes from toxic effects of α-syn fibrils. Elucidation of molecular mechanisms of the observed neuroprotection activity revealed that TMD NFs activate exocytosis and autophagy of toxic protein aggregates. This leads to the suppression of α-syn-induced unfolded protein response (UPR) of the endoplasmic reticulum (ER), and, as a result, a decrease in ROS levels in both neurons and mitochondria. Furthermore, TMD NFs improve mitochondrial health simultaneously facilitating mitochondrial biogenesis. These results demonstrate the therapeutic potential of TMD NFs as a neuroprotective drug candidate capable of decelerating PD.

## METHODS

### Synthesis of TMD NFs

Synthesis of MoS_2_ and MoSe_2_ NFs was conducted using protocols previously described by Mitchell et al.^51^ Fabricated TMD NFs were washed with MilliQ water and ethanol and stored as dry powders at room temperature. Scanning electron microscopy (SEM) at the Material Characterization Facility at Texas A&M University was performed to confirm expected morphological parameters of the nanostructures [RRID: SCR_022202].

### Cell Cultures

Rat dopaminergic midbrain N27 neurons were purchased from American Type Culture Collection (ATCC). Neurons were grown in RPMI 1640 cell medium (Thermo Fisher Scientific) supplemented with 10% heat-inactivated fetal bovine serum (Invitrogen). Rat CTX TNA2 and DI TNC1 astrocyte cells were purchased from ATCC and cultured in Dulbecco’s modified Eagle’s medium (DMEM) supplemented with 10% heat-inactivated fetal bovine serum (Invitrogen). All cell types were cultured in T-75 flasks (Thermo Fisher Scientific) and incubated at 37 °C in a 5% CO_2_ environment. All cell types were utilized prior to the 10^th^ passage to ensure reliability of the results.

### Expression and Purification of α-Syn

To express α-syn, the pET21a-α-synuclein plasmid was transformed into BL21 *Escherichia coli* (*E. coli*) strain Rosetta. The transformed bacteria were incubated at 37 °C in Luria broth (LB) media until reaching OD600 of 0.8-1.2. Next, cells were exposed to 1 mM of IPTG to induce expression of α-syn. Induced bacteria were incubated for additional 4-5 h under the same experimental conditions. Bacteria were collected by centrifugation at 8,000 rpm for 10 min. Bacterial pellets were resuspended in Tris lysis buffer (10 mM EDTA, 150 mM NaCl, 50 mM Tris HCl, pH 7.4) and placed in a water bath pre-heated to 78 °C for 30 min. Next, bacterial suspensions were centrifuged at 16,000 rpm for 40 min to pellet cellular debris that were discarded. Supernatant was collected and treated with streptomycin sulfate (10% solution, 136 μL/mL) in glacial acetic acid (228 μL/mL) followed by centrifugation at 16,000 rpm for 20 min to precipitate bacterial proteins, lipids, and nucleic acids. α-Syn present in the supernatant was precipitated by saturated ammonium sulfate ((NH_4_)_2_SO_4_). The pellet was then resuspended and washed with ammonium acetate (100 mM NH_4_(CH_3_COO)) and absolute ethanol. Harvested α-syn was resuspended in phosphate buffer solution (PBS). Protein samples were concentrated using centrifugation in Amicon Ultra 10 kDa protein filters (Merck Millipore).

Prior to size exclusion chromatography (SEC), protein samples were centrifuged at 14,000 g for 30 min to remove aggregates and impurities. For each SEC cycle, 500 μL of concentrated protein was syringe injected into an AKTA pure FPLC system (GE Healthcare) equipped with a gel-filtration column (Superdex 200 10/300) that was maintained at 4 °C. Proteins were eluted isocratically using PBS, pH 7.4 at a flow rate of 0.5 mL/min. Collected α-syn was reconcentrated in Amicon Ultra 10 kDa protein filters (Merck Millipore) with a final concentration of 400 μM.

### α-Syn Aggregation

A 500 μL solution of α-syn was aggregated in conical microcentrifuge tubes at 37 °C at 510 rpm for 5 days. Fibril formation was confirmed using infrared (IR) spectroscopy and atomic force microscopy (AFM). Mature fibrils were stored at −20 °C until further utilization.

### α-Syn-Induced Cytotoxicity

To model cytotoxic effects of α-syn fibrils, α-syn aggregates were added to neurons and astrocytes. For this, neurons and astrocytes were kept overnight at 37 °C in 5% CO_2_ to reach ∼ 90% confluency. Fibrils were sonicated for 10 min prior to administration to cells. Sonicated α-syn was administered to cells via addition to the media at a final concentration of 40 μM. Cells were allowed to incubate for 12 h with the aggregates before the addition of TMD NFs.

### TMD NFs Treatments

After 12 h induction with α-syn, 10% of the media was removed from each sample. A 2 mg/mL stock of MoS_2_ or MoSe_2_ NFs was resuspended in PBS. The 2 mg/mL stock was probe sonicated for 30 s to completely solubilize the NFs. The 2 mg/mL stock was then serially diluted into the three tested concentrations: 1.0 mg/mL, 0.5 mg/mL, and 0.1 mg/mL. Equivalent volumes of the tested concentrations of NFs were administered to cells. Control cells received equivalent volumes of PBS. The final volume of NFs for each sample is 10% solution consistent across all cell types and prepared concentrations of NFs.

### Flow Cytometry Assays

To analyze ROS levels, cells were stained with CellROX Deep Red Reagent (Thermo Fisher Scientific). To investigate the direct health of mitochondria, cells were stained with MitoProbe JC-1 (Thermo Fisher Scientific). The dyed cells were analyzed using an LSRII BD flow cytometer (BD). Data analysis was conducted using LSRII software.

### Enzyme-linked Immunosorbent Assay (ELISA)

Changes in mitochondrial biogenesis were analyzed using the MitoBiogenesis In-Cell ELISA Colorimetric Kit (Abcam). First, cells were fixed in the plate using BD Cytofix Fixation Buffer (BD Biosciences). Cells were permeabilized, blocked, and incubated with primary antibodies. Cells were washed and administered secondary antibodies. After washing, cells were treated with AP development solution. The absorbance (λ=405 nm) of each well was measured using a Tecan plate reader (Tecan Life Sciences) to quantify expression of succinate dehydrogenase subunit A (SDH-A). The media was removed, and the cells were administered horseradish peroxidase (HRP) development solution. Absorbance (λ=600 nm) was measured using a Tecan plate reader (Tecan Life Sciences) to quantify expression of cytochrome c oxidase subunit I (COX-I).

### Quantitative Polymerase Chain Reaction (qPCR)

All primers were designed based on rat gene sequences documented in the NCBI database and developed using the SnapGene custom oligonucleotide PCR primer generation software, Table 1. All primers were fabricated by Integrated DNA Technologies (IDT). RNA was extracted from the cell pellet using a GeneJET RNA Purification Kit (Thermo Scientific). RNA concentration was measured using a NanoDrop One instrument. Next, RNA was converted to complimentary DNA (cDNA) using SuperScript II Reverse Transcriptase (Invitrogen) with random primers (Invitrogen). Synthesized cDNA, constructed primers, and Luna Universal qPCR Master Mix (New England Biolabs) were mixed and placed in a QuantStudio 7 Flex Real-Time PCR System (Thermo Scientific). Changes in gene expression levels were calculated using the comparative C_T_ method (2^-ΔΔC_T_). GAPDH was used as a reference gene to calculate 2^-ΔΔC_T_.

**Table 1.**
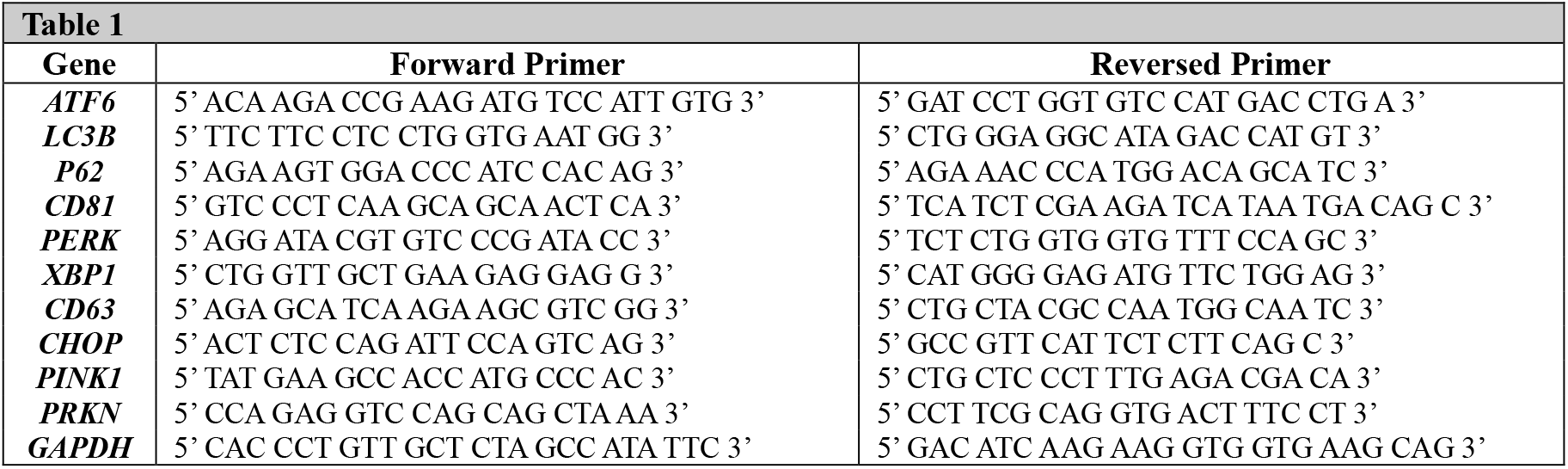
Sequence of forward and reversed primers used for qPCR.

### Stress Granule Imaging

After washing with PBS, cells were fixed with 100 μL BD Cytofix™ Fixation Buffer (BD Biosciences). Next, cells membranes were permeabilized with Triton-X and blocked with bovine serum albumin (BSA) to limit unspecific protein binding. Administration of T-cell intracellular antigen 1-related protein (TIA-R) fluorescent primary antibodies was conducted at 4 °C overnight to ensure complete binding. Treated cells were washed to remove unbound fluorescent antibodies. Cells were imaged using an EVOS M5000 microscope (Thermo Fisher Scientific) to track the presence of induced stress granules.

### C. elegans assays

Transgenic fluorescently tagged alpha synuclein strain NL5901, with genotype pkIs2386 [unc-54p::alphasynuclein::YFP + unc-119(+)], was purchased from University of Minnesota Caenorhabditis Genetic Center (CGC). *C. elegans* were kept on nematode growth medium (NGM) plates seeded with OP50 *E. coli* and at 20 °C until reaching an egg-producing age. To ensure accurate life span tracking, age synchronization was conducted by collecting all worms and eggs and bleaching the solution to remove all adult worms, according to Sutphin & Kaeberlein’s protocol ^55^. Three days after synchronization, nematodes were transferred onto experimental NGM plates doped with fluorodeoxyuridine (FUDR) to prevent eggs from hatching. A total of ten nematodes were transferred to each experimental plate. The plates were seeded with *E. coli* supplemented with different concentrations of TMD NFs. NF supplementation was conducted by mixing concentrated 2% stocks with 10x concentrated OP50 E. coli at a 1:1 ratio before plating, quickly drying, and UV irradiating for a final concentration of 1% TMD NFs. Daily counts of the number of alive and dead nematodes were conducted to generate a Kaplan-Meier survival curve and calculate statistical analyses.

### Fluorescent Imaging of C. elegans

Fluorescent imaging of endogenous α-syn puncta was conducted via an EVOS M5000 microscope (Thermo Fisher Scientific). Individual nematodes were placed in 5 μL of 50 mM sodium azide (NaN_3_) for immobilization. The nematodes were imaged in the green fluorescent protein (GFP) light channel. Five nematodes were imaged for each experimental condition at each given time point. Images were analyzed using ImageJ software to quantify the number of fluorescent puncta present in each worm. The average number of puncta across the five nematodes was recorded.

## RESULTS

### TMD NF Treatments Alleviate α-Syn-Induced Mitochondrial Impairment and Boost Mitochondrial Biogenesis in Neurons and Astrocytes

Endocytosis of α-syn aggregates by neurons and astrocytes triggers a cascade of pathological processes that results in a depolarization of mitochondrial membranes, and consequently, impaired ATP biosynthesis, which ultimately leads to a cell death ^56-59^. Expanding upon this, JC-1 assay was used to determine the extent to which TMD NFs could reverse α-syn-induced mitochondrial membrane potential. An uptake of α-syn fibrils induced significant mitochondrial damage in both neurons and astrocytes. Compared to the control, N27 neurons experienced a 257% increase, while DI astrocytes a 327% increase in mitochondrial damage upon exposure to α-syn aggregates. Similarly to N27 neurons, α-syn fibrils triggered a 251% increase in the mitochondrial damage in CTX astrocytes [**Figure 1**].

**Figure 1:**
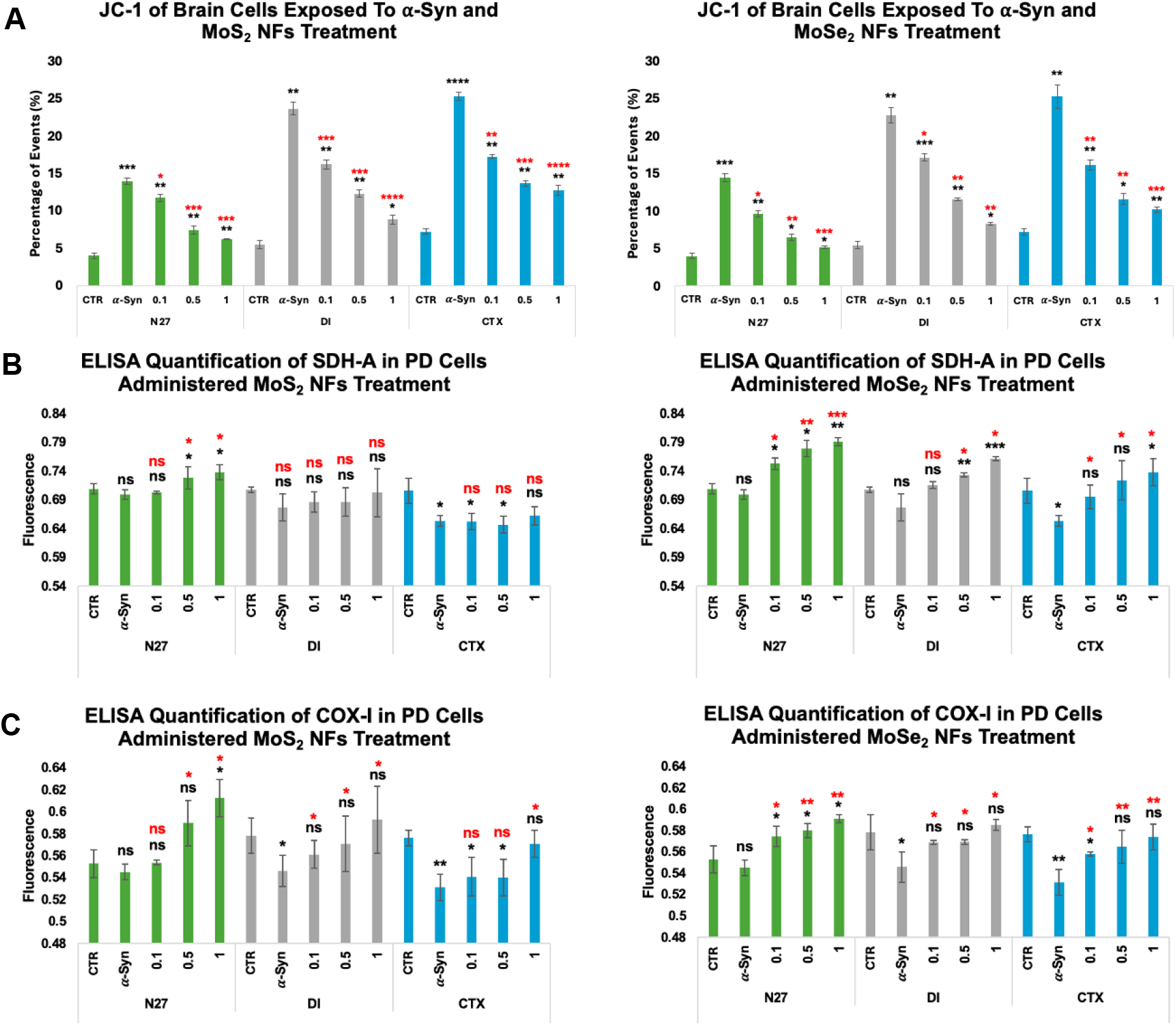
**[A]** JC-1 mitochondrial damage assay conducted on N27 neurons (green), DI astrocytes (gray), and CTX astrocytes (blue) treated with MoS_2_ NFs [left] and MoSe_2_ NFs [right]. Mitochondrial biogenesis ELISA quantifications of nuclear encoded SDH-A **[B]** and mitochondrially encoded COX-I [**C**] when treated with MoS_2_ NFs [left] and MoSe_2_ NFs [right] in N27 neurons (green), DI astrocytes (gray), and CTX astrocytes (blue). Experimental groups of cells receiving no α-syn or NF treatment (CTR), cells only exposed to α-syn (α-Syn), and cells exposed to α-syn and subsequent NF treatments of 0.1 mg/mL (0.1), 0.5 mg/mL (0.5), or 1.0 mg/mL (1). Student’s t-test was utilized to analyze statistical significance of reported results, with * P< 0.05, ** P< 0.005, *** P< 0.0005, and NS – not significant. Black significance markings relate to comparisons made to control cells that did not receive α-syn or NF treatments. Red significance markings relate to comparisons made to PD-induced control cells that were exposed to α-syn but no TMD NF treatments.

The subsequent (12h) treatment of these neurons with different concentrations of MoS_2_ NFs resulted in 15.8% to 55.4% decreases in the magnitude of α-syn-induced mitochondrial damage. DI and CTX astrocytes treated with MoS_2_ NFs also experienced a strong dose-dependent therapeutic relief, which resulted in 31.5% to 62.7% (DI) and 31.9% to 49.7% (CTX) decrease in the mitochondrial impairment. Similar effects were observed for MoSe_2_ NFs in both neurons and astrocytes. N27 neurons exposed to α-syn fibrils and subsequently (12h) treated with MoSe_2_ NFs exhibited 33.5% to 64.0% reduction in the mitochondrial impairment. In DI and CTX astrocytes exposed to α-syn fibrils, MoSe_2_ NFs caused a strong concentration-dependent reduction in mitochondrial damage ranging from 24.9% to 63.7%. These results indicate that both MoSe_2_ and MoS_2_ NFs efficiently rescue cell mitochondrial from cytotoxic effects caused by α-syn aggregates.

It is important to determine whether TMD NFs only rescue cell mitochondria or also facilitate the synthesis of novel mitochondria, a process known as mitochondrial biogenesis.^60-62^. To answer this question, changes in the expression of two mitochondrial proteins, succinate dehydrogenase subunit A (SDH-A) and cytochrome c oxidase subunit I (COX-I) were analyzed. ELISA revealed that exposition of neurons and astrocytes to α-syn fibrils caused a strong downregulation in the expression of both SDH-A and COX-I [**Figure 1**]. N27 neurons experienced only a small, 1.34% and 1.39% decrease in the expression of SDH-A and COX-I protein levels, respectively. However, a substantially higher magnitude (4.32% for SDH-A and 5.60% for COX-I) of downregulation of mitochondrial biogenesis was observed in DI, as well as in CTX astrocytes (7.48% for SDH-A and 7.82% for COX-I). These results indicate that endocytosis of α-syn aggregates strongly suppresses mitochondrial biogenesis in both neurons and astrocytes.

A dose-dependent upregulation in mitochondrial biogenesis was observed in N27 neurons after administration of MoS_2_ and MoSe_2_ NF. As the concentration of MoS_2_ NFs increased from 0.1 to 1.0 mg/mL, SDH-A was upregulated from 0.51% to 5.49%. At the same concentrations of MoS_2_, COX-I was upregulated from 1.57% to 12.35%. When treated with MoSe_2_ NFs, the expression of SDH-A was upregulated from 7.63% to 13.11%, while COX-I expression was upregulated from 5.38% to 8.39%. Similarly to neurons, DI astrocytes underwent a dose-dependent upregulation of mitochondrial biogenesis upon receiving TMD NF subsequently to α-syn aggregates. As the concentration of MoS_2_ NFs increased from 0.1 to 1.0 mg/mL, an upregulation of SDH-A from 1.40% to 3.74% was observed. COX-I upregulation was found to increase from 2.76% to 8.60% in the cells exposed to 0.1 to 1.0 mg/mL of MoS_2_ NFs. When treated with MoSe_2_ NFs, SDH-A experienced an upregulation from 5.68% to 12.43%, while COX-I expression was alleviated from 4.16% to 7.22%. CTX astrocytes treated with MoS_2_ NFs had little to no change in the levels of SDH-A expression. At the highest concentration of MoS_2_ NFs, only a 1.37% increase in SDH-A expression was observed. At the same time, a significant upregulation (1.81% to 7.43%) of COX-1 expression was observed in CTX astrocytes exposed to MoS_2_ NFs. CTX astrocytes treated with MoSe_2_ NFs showed a more prominent dose-dependent response for both mitochondrial proteins. The expression of SDH-A was elevated by 6.44%, 10.80%, and 13.02%. The COX-I protein experienced an increase in the expression from 5.09% to 8.03% as the concentration of MoSe_2_ NFs increased. These results indicate that MoS_2_ and MoSe_2_ NFs not only protect mitochondria from α-syn aggregates but also boost mitochondrial biogenesis in both neurons and astrocytes.

### TMD NF Treatments Alleviate α-Syn-Induced ROS and Reverse Cellular Stress Granules

Internalization of aggregated α-syn by neuronal cells results in an increase in ROS^63, 64^ and translation of stress granule proteins from the nucleus to the cytosol^65^. Flow cytometry-based ROS assay was used to determine the extent to which TMD NFs altered ROS levels in neurons and astrocytes. An exposition of N27 neurons, DI, and CTX astrocytes to α-syn aggregates caused a significant increase in the ROS levels. Specifically, N27 neurons experienced a 213% increase, while DI and CTX astrocytes had 132% and 59.3% increase in ROS, respectively [**Figure 2**]. A subsequent (12h) treatment of neurons and astrocytes exposed to α-syn aggregates with MoS_2_ NFs resulted in a dose-dependent decrease in the cellular ROS levels. N27 neurons exposed to the lowest concentration of MoS_2_ NFs (0.1 mg/mL) experienced a moderate (13.4%) ROS reduction. Higher concentrations of MoS_2_ NFs (0.5 mg/mL and 1.0 mg/mL) caused a substantially higher (24.8% and 43.8%, respectively) decrease in the ROS levels. Similar results were observed for DI astrocytes. The lowest concentration of MoS_2_ NFs resulted in 7.6% reduction, while administration of 0.5 mg/mL and 1.0 mg/mL MoS_2_ NFs led to the significantly larger decrease (31.5% and 41.8%, respectively) in ROS levels. Similar magnitudes of MoS_2_ NFs-triggered decrease in ROS levels in CTX astrocytes were also observed. Specifically, a decrease in ROS to 4.9%, 23.6%, and 36.1% was observed in the cells exposed to 0.1, 0.5, and 1.0 mg/mL MoS_2_ NFs.

**Figure 2:**
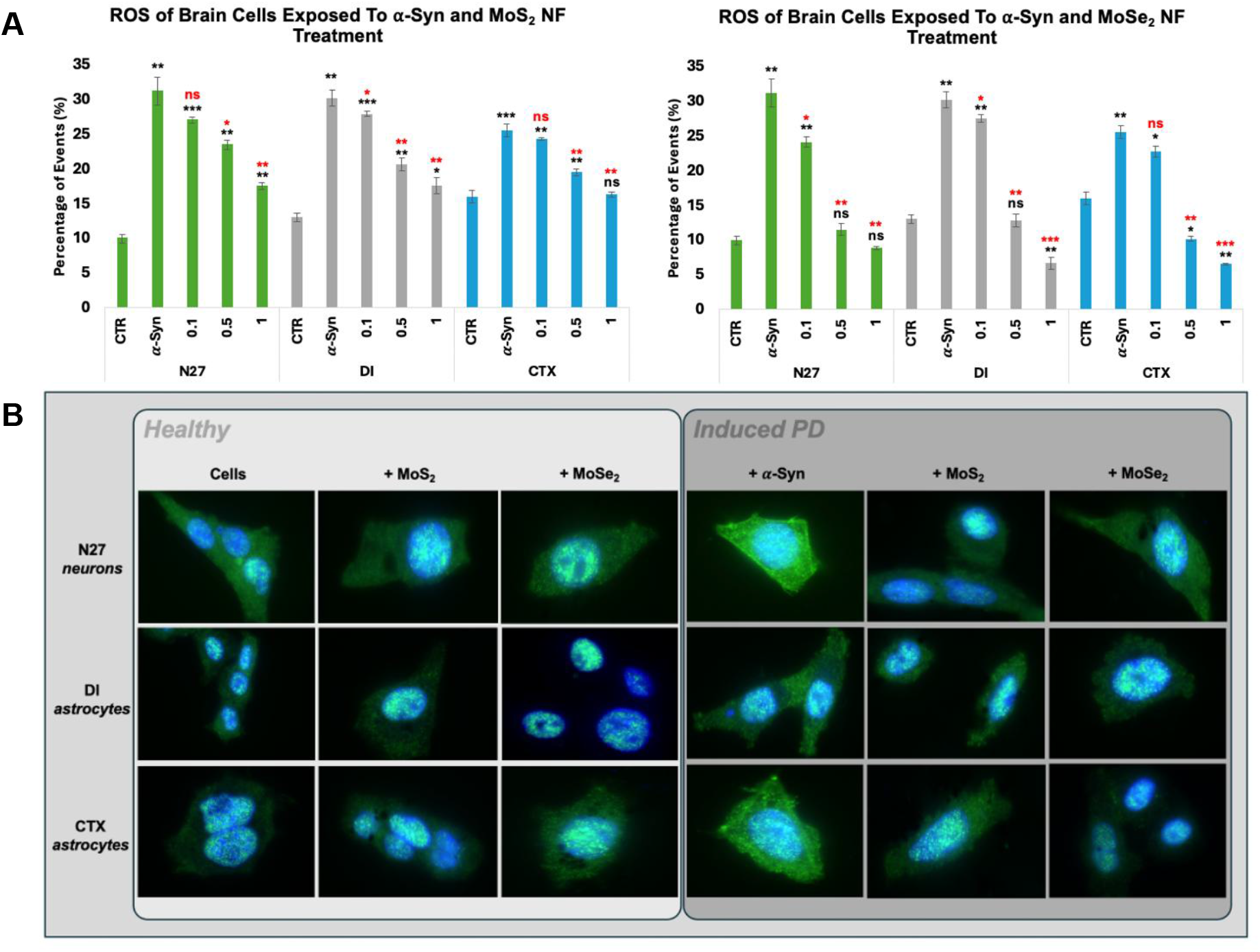
**[A]** ROS assay conducted on N27 neurons (green), DI astrocytes (gray), and CTX astrocytes (blue) treated with MoS_2_ NFs [left] and MoSe_2_ NFs [right]. Experimental groups of cells receiving no α-syn or NF treatment (CTR), cells only exposed to α-syn (α-Syn), and cells exposed to α-syn and subsequent NF treatments of 0.1 mg/mL (0.1), 0.5 mg/mL (0.5), or 1.0 mg/mL (1). Student’s t-test was utilized to analyze statistical significance of reported results, with * P< 0.05, ** P< 0.005, *** P< 0.0005, and NS – not significant. Black significance markings relate to comparisons made to control cells that did not receive α-syn or NF treatments. Red significance markings relate to comparisons made to PD-induced control cells that received α-syn but no TMD NF treatments. [**B]** Immunofluorescent imaging of the stress granule protein TIA-R in healthy [light gray box] N27 neurons (top row), DI astrocytes (middle row), and CTX astrocytes (bottom row) when treated with no NFs (first column), MoS_2_ NFs (second column), or MoSe_2_ NFs (third column). Images of induced PD [dark gray box] N27 neurons (top row), DI astrocytes (middle row), and CTX astrocytes (bottom row) when treated with no NFs (fourth column), MoS_2_ NFs (fifth column), or MoSe_2_ NFs (sixth column). Cell images taken at 400X magnification.

A much larger magnitude in the suppression of ROS levels was observed in neurons and astrocytes exposed to MoSe_2_ NF. Specifically, N27 neurons exposed to 0.1 mg/mL MoSe_2_ NFs experienced a 22.5% reduction in ROS levels. At 0.5 and 1.0 mg/mL, the reduction of ROS in 63.0% and 71.6% were recorded. A similar magnitude in the reduction of ROS was observed in DI astrocytes exposed to MoSe_2_ NFs. The lowest concentration of NFs resulted in 8.9% reduction of ROS. At the same time, medium and high (0.5 and 1.0 mg/mL) concentrations of MoSe_2_ NFs caused a suppression of ROS levels on 57.6% and 77.8%, respectively. A graduate decrease in ROS levels (11.1%, 60.3%, and 74.3%) in CTX astrocytes was also observed with a graduate increase in the concentration of MoS_2_ NFs (0.1, 0.5 and 1.0 mg/mL). It should be noted that for all cell types, the highest concentration of NFs reduced ROS levels to the baseline observed in the cells experienced no α-syn. These results show that both MoSe_2_ and MoS_2_ NFs drastically reduce α-syn-induced ROS levels in neurons and astrocytes.

Cells can form, maintain, and dissociate dynamic condensates known as stress granules^66^. These condensates are comprised of RNA and RNA-binding proteins. Stress granules regulate multiple cellular processes focused on the minimization of cellular stress^66, 67^. In order to probe the effects TMD NF therapeutics have on cellular stress mechanisms, immunofluorescent tracking of T-cell intracellular antigen 1-related (TIA-R) protein was conducted. TIA-R is a nucleic acid binding protein localized inside the nucleus under normal functioning conditions but vacates to the cytosol upon activation via cellular stress in pathological environments. To ensure that MoS_2_ and MoSe_2_ NFs caused no cellular stress, neurons and astrocytes were treated with TMD NFs. Immunofluorescent staining of these cells revealed confined localization of TIA-R in the nuclei, indicating that TMD NF alone did not induce any stress in neurons and astrocytes [**Figure 2**]. Immunofluorescent staining revealed the appearance of stress granules in the cytosol of neurons and astrocytes exposed to α-syn aggregates. From all analyzed cells, neurons had the most drastic response to α-syn aggregates with nearly complete evacuation of TIA-R from the nuclei. CTX astrocytes were also observed to form high levels of stress granules in the cytosol. DI astrocytes, however, did not respond with the same degree of magnitude, forming fewer stress granules in response to α-syn aggregates.

All cell types treated with MoS_2_ and MoSe_2_ NFs experienced a significant reduction in stress granule formation. It was found that TMD NFs administration resulted in the re-localization of TIA-R back to the nuclei, with very few fluorescent signals still present in the cytosol. Cells receiving MoSe_2_ NFs treatments experienced even greater stress relief compared to their MoS_2_ NFs counterparts. Thus, TIA-R imaging confirms that TMD NFs do provide neuroprotection via regulating stress response, but more in-depth investigation is needed to better determine the complete molecular mechanistic machinery.

### Elucidation of Neuroprotective Mechanisms of TMD NF

Accumulation of α-syn aggregates in neuronal cells triggers the unfolded protein response (UPR)^68, 69^ in the endoplasmic reticulum (ER)^70, 71^, reduction of exocytosis^72^, and increased mitophagy^58, 73^, [**Figure 3**]. Using quantitative polymerase chain reaction (qPCR), changes in the expression of genes involved in UPR, exocytosis, and mitophagy were determined in neurons and astrocytes exposed to α-syn fibrils followed by TMD NFs treatment, as well as α-syn fibrils alone (control).

**Figure 3:**
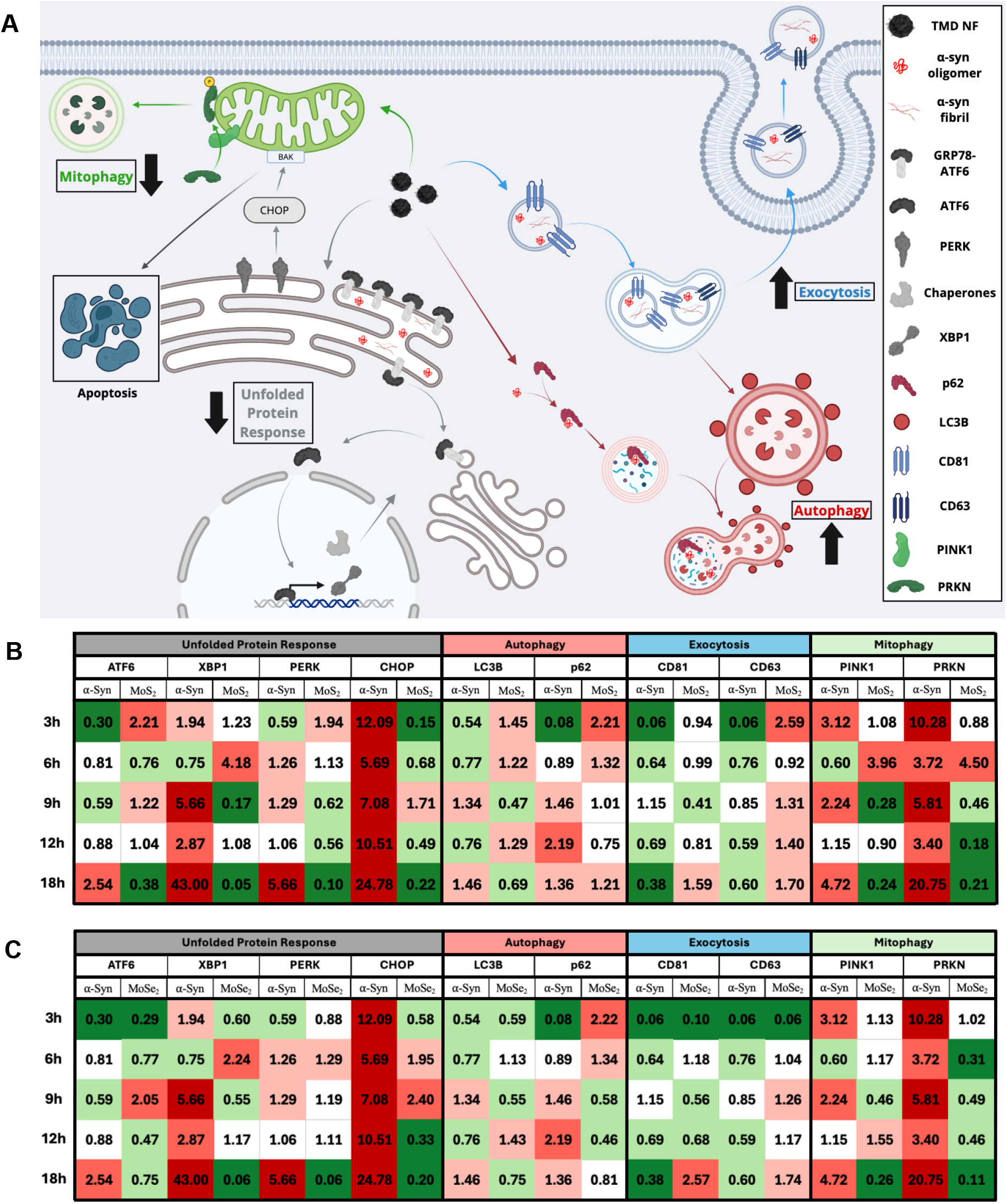
**[A]** Schematic of known cellular pathways negatively affected by introduction of aggregated α-syn, with the four major cascades color coded: unfolded protein response (gray), autophagy (red), exocytosis (blue), and mitophagy (green). Heat map of calculated changes in gene expression via qPCR for N27 neurons treated with MoS_2_ NFs [**B**] or MoSe_2_ NFs [**C**] at 3h, 6h, 9h, 12h, and 18h post NF treatment and compared to control cells. The 2^−ΔΔ*CT*^values are shaded as: 0.01-0.40 (dark green), 0.41-0.80 (light green), 0.81-1.20 (white), 1.21-2.0 (light red), 2.1-5.0 (bright red), 5.0+ (maroon).

Impregnation of aggregated α-syn into neuronal cells significantly increases the UPR via upregulation of key genes, including protein kinase R-like endoplasmic reticulum kinase (PERK), activating transcription factor 6 (ATF6), C/EBP homologous protein (CHOP), and X-box binding protein 1 (XBP1). Administration of both MoS_2_ and MoSe_2_ NF alleviated the pathogenic burden of aggregated α-syn in N27 neurons on the level of genetic regulation, [**Figure 3**]. Specifically, MoS_2_ NFs downregulated all four UPR genes after 18 hours of TMD NFs administration. ATF6 experienced a 62% reduction, while XBP1 underwent a 95% downregulation in the cells exposed to α-syn fibrils followed by MoS_2_ NFs treatment. PERK and CHOP each experienced major downregulation in such cells, with 90% and 78% reductions, respectively. Administration of MoSe_2_ NFs resulted in downregulation in all four chosen UPR genes at the 18h timepoint. ATF6 experienced a 25% reduction, having the least impact. The other UPR genes, however, had very strong downregulations with 94%, 94%, and 80% reductions for XBP1, PERK, and CHOP, respectively.

In neurons, accumulation of aggregated proteins also first triggers downregulation of autophagy genes, microtubule-associated protein 1 light chain 3 beta (LC3B) and sequestosome 1 (p62). This initial suppression is followed by upregulation of these genes as neurons degrade and autophagy amyloid aggregates. MoS_2_ NFs upregulated both LC3B and p62 after 3h and 6h of treatment, indicating their innate ability to advance cell autophagy and quickly mitigate amyloid-induced stress in the cells. LC3B was found to be subsequently downregulated at the 18h, while p62 maintained upregulated. The exocytosis response was upregulated at the later timepoints, indicating an TMD-facilitated ability to remove α-syn fibrils. MoSe_2_ NFs triggered upregulation of p62 2.22-fold at the 3h timepoint and then tapers off for the remaining timepoints as the presence of aggregates decreases.

Cluster of differentiation 81 and 63 (CD81 and CD63) are exosomal markers that can be used to quantify changes in the cell exocytosis activity. Both CD81 and CD63 were strongly downregulated in the cells exposed to α-syn fibrils alone. In the neurons exposed to α-syn fibrils followed by TMD NFs treatment, CD81 was unchanged in the two early time points and then experienced a downregulation at the 9h timepoint. From that point forward, there was a great upregulation, hinting at the cell’s ability to exocytose increasing. CD63 was initially upregulated at the 3h timepoint and remained upregulated. More research will be conducted to identify and quantify the contents of the exosomes to provide more insight into cellular response. In the cells exposed to MoSe_2_ NFs, CD81 was downregulated at the early timepoints, then experienced an upregulation up to 2.57x at 18h. CD63 was upregulated much faster, with upregulation occurring at the 9h, 12h, and 18h timepoints, with a maximum upregulation of 1.74x at the 18h timepoint.

As was discussed above, amyloid aggregates damage cell mitochondria, which in turn, results in the upregulation of mitophagy. Two major players involved in mitophagy are the PTEN-induced putative kinase 1 (PINK1) and parkin (PRKN) proteins. qPCR revealed that neurons exposed to α-syn fibrils alone experienced strong upregulation of both Pink1 and Prkn. MoS_2_ NFs strongly affected the regulatory elements of the mitophagy pathway. At the 9h, 12h, and 18h timepoints, both PINK1 and PRKN were significantly downregulated, indicating the cells decreased need for mitophagy due to the presence of healthier mitochondria. Administration of MoSe_2_ NF also led to downregulation of mitophagy. At the 18h timepoint, PINK1 experienced a 74% reduction. The PRKN gene experienced a reduction at the 6h, 9h, 12h, and 18h timepoints, culminating at an 89% reduction for the 18h timepoint. Summarizing, both MoS_2_ and MoSe_2_ NF were able to downregulate the UPR and mitophagy responses, while upregulating autophagy and exocytosis pathways in a neuroprotective manner.

### Elucidation of Neuroprotective Effects of TMD NFs In Vivo

*C. elegans* are a well-characterized and established multicellular organism utilized for a multitude of experimentations, including drug screenings^74-76^, understanding mechanisms of neurodegenerative diseases^77-79^, and fundamental biology^80, 81^. To probe the neuroprotective effects that TMD NFs, *C. elegans* that overexpress the α-syn protein were utilized. The overexpressed α-syn was tagged with green fluorescent protein allowing amyloid aggregates to form puncta and be tracked using fluorescent microscopy. MoS_2_ or MoSe_2_ NFs were supplemented in the nematodes’ diet. The quantification of fluorescent puncta can be used to track the rate of α-syn accumulation in nematodes. Additionally, *C. elegans* lifespan was analyzed for further determination of TMD NF neuroprotective properties.

*C. elegans* receiving MoS_2_ NFs in their diet exhibited a slight increase in life expectancy compared to the control group. The control nematodes had a calculated survival expectancy (*p*=50%) of 18.5 days, as shown in **Figure 4**. *C. elegans* supplied with 0.1 mg/mL MoS_2_ NFs in their diet did not experience a shift in the *p*=50% (18.5 days). While nematodes receiving 0.5 mg/mL supplementation of the same nanostructures had an increase in p=50% of one day (19.5 days), the nematodes receiving the highest concentration of MoS_2_ NFs (1.0 mg/mL) demonstrated an extension of p=50% of two days (20.5 days).

**Figure 4:**
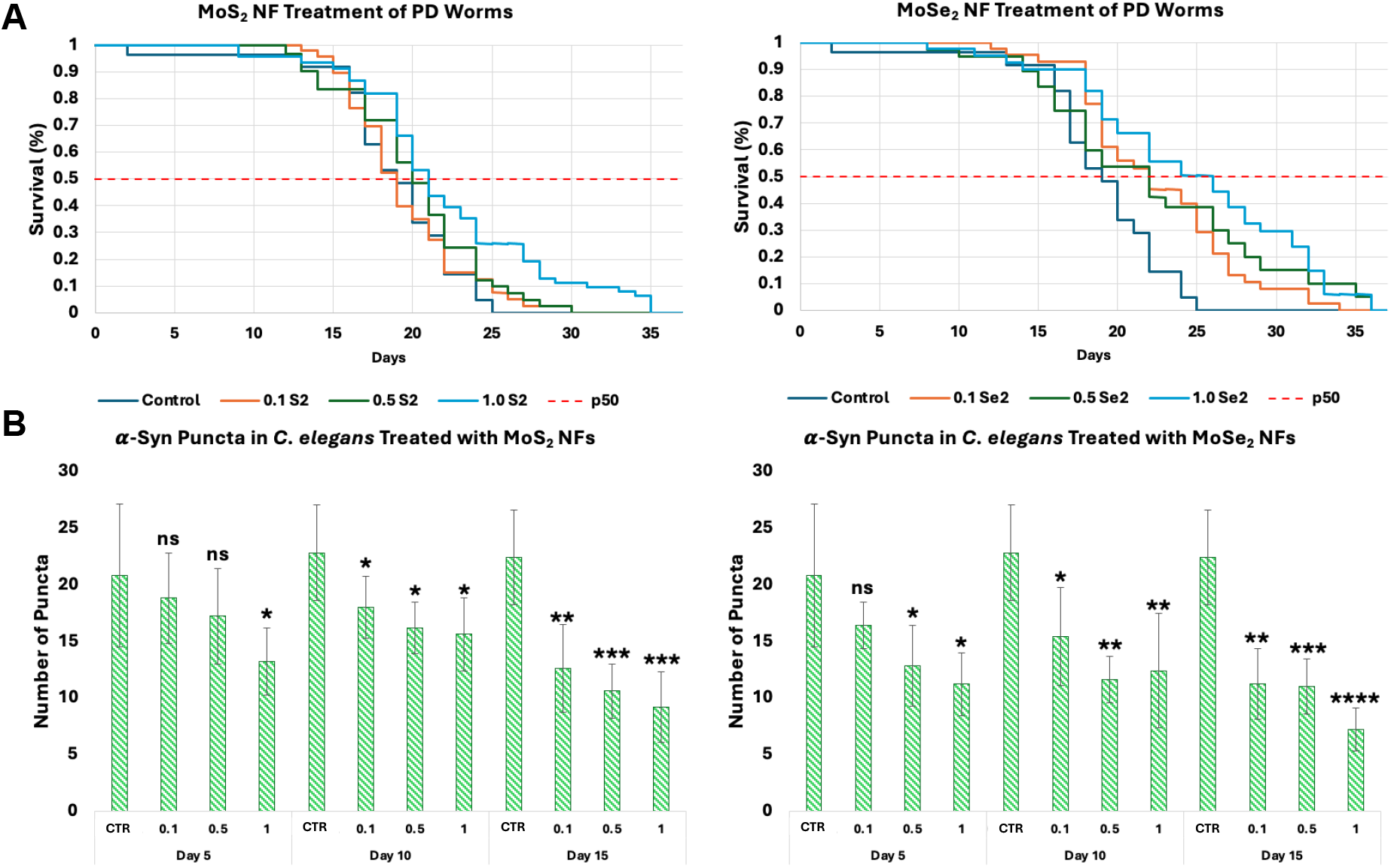
[**A]** Calculated Kaplan-Meier life expectancy curves of PD *C. elegans* when administered MoS_2_ NFs [left] or MoSe_2_ NFs [right] at 0.1 mg/mL (orange), 0.5 mg/mL (green), 1.0 mg/mL (light blue) or 0 mg/mL (dark blue) concentrations. [**B]** Image analyzed fluorescent α-syn aggregates in the pharynx of PD *C. elegans* when treated with MoS_2_ NFs [left] or MoSe_2_ [right] after 5, 10, or 15 days of treatment. Experimental groups of PD nematodes receiving no NF treatment (CTR), or administered NF treatments of 0.1 mg/mL (0.1), 0.5 mg/mL (0.5), or 1.0 mg/mL (1). Student’s t-test was utilized to analyze statistical significance of reported results, with * P< 0.05, ** P< 0.005, *** P< 0.0005, **** P< 0.00005 and NS – not significant. Significance markings coincide with comparisons made to each respective timepoint’s control nematodes that did not receive any NF treatment.

Upon investigation of the quantity of puncta present in each group of worms on Day 5, Day 10, and Day 15, a clear dose-dependent reduction in the accumulation rate of α-syn was clear, [**Figure 4**]. On Day 5, control nematodes had an average of 20.8 counted puncta (20.8 ±6.34). As the concentration of MoS_2_ increased, there was a reduction of 9.62%, 17.31%, and 36.54% when administered 0.1 mg/mL (18.8 ±3.96), 0.5 mg/mL (17.2 ±4.21), and 1.0 mg/mL (13.2 ±2.95), respectively. On Day 10, the control group had an average of 22.8 puncta present (22.8 ±4.21). A dose-dependent reduction of 21.05%, 28.95%, and 31.58% was observed for the 0.1 mg/mL (18.0 ±2.74), 0.5 mg/mL (16.2 ±2.28), and 1.0 mg/mL (15.6 ±3.21) groups, respectively. The greatest reductions in puncta quantity were observed on Day 15. The control group had 22.4 ±4.16 puncta count. When administered 0.1 mg/mL MoS_2_, there was a 43.75% reduction in the puncta count (12.6 ±3.85). Nematodes supplemented with 0.5 mg/mL had a reduction of 52.68% (10.6 ±2.41) in puncta present. The greatest reduction was observed in nematodes receiving the highest concentration (1.0 mg/mL) of MoS_2_ NFs, with puncta count reduced by 58.93% (9.2 ±3.11). Although exact molecular mechanisms of MoS_2_ NF triggered amyloid reduction remains unclear, these results help explain the dose-dependent increase in life expectancy of *C. elegans*.

MoSe_2_ NFs exhibited a significant increase in life expectancy when administered to *C. elegans*. Nematodes supplied with 0.1 mg/mL and 0.5 mg/mL both experienced an increase in *p*=50% of three days (21.5 days). This represents a 16.22% increase in life expectancy. The nematodes receiving the highest dosage of MoSe_2_ (1.0 mg/mL) experienced the strongest magnitude of extension in life expectancy. This group demonstrated a seven-day extension in life expectancy, with a *p*=50% value of 25.5 days.

Analysis of fluorescent puncta in the nematodes receiving MoSe_2_ NF treatments revealed a significant reduction in amyloid burden. On Day 5, a clear dose-dependent reduction in puncta was observed. As the MoSe_2_ NF concentrations increased from 0.1 to 1.0 mg/mL, there were reductions of 21.15%, 38.46%, 46.15% calculated (16.4 ±2.07, 12.8 ±3.56, 11.2 ±2.77), respectively. At the next timepoint, Day 10, *C. elegans* that received MoSe_2_ NFs continued to exhibit a drastic decrease in the number of puncta. There were decreases of 32.46%, 49.12%, and 45.61% as the concentration of NFs increased from 0.1 mg/mL (15.4 ±4.34) to 0.5 mg/mL (11.6 ±2.07) and 1.0 mg/mL (12.4 ±5.03). Day 15 revealed the most drastic reduction of puncta in nematodes administered MoSe_2_ NF treatment. Upon receiving 0.1 mg/mL MoSe_2_ NFs, there was a 50.0% reduction in the counted puncta (11.2 ±3.11). The nematodes treated with 0.5 mg/mL MoSe_2_ experienced a 50.89% reduction in aggregates (11.0 ±2.45). The highest magnitude of reduction was found in *C. elegans* exposed to 1.0 mg/mL MoSe_2_ NFs, with a 67.86% decrease in the puncta present at Day 15 (7.2 ±1.92). These results revealed that MoSe_2_ and MoS_2_ NFs exert similar therapeutic effects, however, MoSe_2_ enacts significantly higher magnitudes of neuroprotective properties in *C. elegans*.

## DISCUSSION

The onset and progression of PD is driven by the pathological aggregation of α-syn and propagation of cytotoxic aggregates from cell to cell^24, 82, 83^. Upon endocytosis, α-syn aggregates damage mitochondria^84, 85^ and ER^86^ which results in increased ROS^87^, stress granule formation^88^, and reduced autophagy^89^, ultimately leading to cell death^90^. Expanding upon this, NF ability to reverse key pathological processes and alleviate cell stress was investigated.

JC-1 assay revealed that TMD NFs prevent α-syn-triggered depolarization of cell mitochondrial. Furthermore, ELISA-based analysis of the expression of COX-I and SDH-A proteins indicated that both MoS_2_ or MoSe_2_ NFs induced mitochondrial biogenesis in neurons and astrocytes. Previously, our group demonstrated that TMD NFs-induced mitochondrial biogenesis was activated via upregulation of the expression of peroxisome proliferator-activated receptor-gamma coactivator (PGC-1α), which in turn, upregulated nuclear respiratory factor (NRFs), the key player in mitochondrial biogenesis^51^. It is important to note that upregulation of PGC-1α was linked to the downregulation of steroid receptor coactivator (SRC-3) and general control non-depressible 5 (GCN5) factors that sense energy excess in the cell. Summarizing, the results of the current study indicate that TMD NFs prevent α-syn-triggered mitochondrial damage simultaneously facilitating mitochondrial biogenesis in neurons and astrocytes. These conclusions are further supported by downregulation of mitophagy in the cells sequentially exposed to α-syn fibrils and TMD NFs.

Although the full molecular mechanism of TMD NFs-suppressed ROS levels in cells is yet to be determined, previously reported findings showed that MoS_2_ or MoSe_2_ NFs exhibited catalase, peroxidase, and superoxide dismutase activities^50^. Thus, TMD NFs are capable of degrading free radicals and remove hydrogen peroxide from the cells minimizing detrimental effect of such toxic species on both neurons and astrocytes. It has been previously shown that an upregulation of PGC-1α activated the expression of steroid receptor coactivator-3 (SIRT-3) and estrogen-related receptor alpha (ERRα) that suppress ROS levels in cells ^91^. Currently reported experimental results demonstrate that MoS_2_ or MoSe_2_ NFs also suppressed UPR of ER which further decreased the magnitude of ROS in both neurons and astrocytes. This evidence is further supported by re-localization of stress granules in the cells sequentially exposed to α-syn fibrils and TMD NFs back to the nuclei. qPCR-based analyses of LC3B, p62, CD81, and CD63, suggest that a decrease in the cell ROS levels could be linked to TMD NFs-facilitated autophagy and exocytosis of toxic amyloid aggregates by neurons and astrocytes.

Utilization of *C. elegans* as a model living system confirmed observed *in vitro* neuroprotective properties of TMD NFs. Fluorescence imaging revealed a suppression of puncta formations in the nematodes exposed to both MoS_2_ or MoSe_2_ NFs. As expected, such amyloid-depleted nematodes exhibited much longer lifespan compared to the control group of *C. elegans*. It was hypothesized that TMD NFs could directly inhibit α-syn overexpressed by *C. elegans* used in the current study. To test this hypothesis, thioflavin T assay was used. *In vitro* aggregation of α-syn in the presence of different concentrations of MoS_2_ or MoSe_2_ NFs did not reveal any inhibition properties of TMD NFs (Figure S1). These results indicate that autophagy and other molecular mechanisms are responsible for the observed reduction of puncta formations in the nematodes exposed to both MoS_2_ or MoSe_2_ NFs.

It is important to note that both *in vivo* and *in vitro* assays used in the current study revealed dissimilar magnitude in the neuroprotective activity of MoS_2_ or MoSe_2_ NFs present at the same concentrations. Specifically, MoSe_2_ NF exhibited a much more potent neuroprotective ability than MoS_2_ NFs. Such difference could be linked to the different chemical structures of the nanomaterials, as well as their size and shape. Detailed elucidation of the relationship between neuroprotection activity, chemical and physical properties of TMD NFs is the subject of a separate study.

## CONCLUSION

Experimental findings reported in the current study demonstrate that MoS_2_ or MoSe_2_ NFs were capable of rescuing both neurons and astrocytes from the cytotoxic effects of α-syn fibrils. The mechanism of neuroprotective activity includes suppression of α-syn-induced UPR in the ER, and TMD NFs-facilitated autophagy and exocytosis of α-syn fibrils. In parallel, MoS_2_ or MoSe_2_ NFs reversed α-syn-induced damage of cell mitochondria while simultaneously stimulating mitochondrial biogenesis via activation of PGC-1α. This resulted in a suppression of mitophagy and a drastic reduction in ROS levels. Utilization of *C. elegans* that overexpressed α-syn confirmed *in vitro* observed dose-dependent effect of TMD NFs. Such nematodes experienced a drastic reduction in the α-syn aggregation which resulted in a drastic increase in *C. elegans* lifespan. Cumulative results suggest TMD NFs are potential neuroprotective therapeutics for PD, with further in-depth studies needed.

### Notes

The authors declare no competing financial interests

## Acknowledgements

We are grateful to the National Institute of Health for the provided financial support (R35GM142869).

## Notes

### Competing Interest Statement

The authors have declared no competing interest.

## References

1. D. Su, Y. Cui, C. He, P. Yin, R. Bai, J. Zhu, J. S. T. Lam, J. Zhang, R. Yan, X. Zheng, J. Wu, D. Zhao, A. Wang, M. Zhou and T. Feng, Projections for prevalence of Parkinson’s disease and its driving factors in 195 countries and territories to 2050: modelling study of Global Burden of Disease Study 2021, BMJ, 2025, 388, e080952.

2. P. Chopade, N. Chopade, Z. Zhao, S. Mitragotri, R. Liao and V. Chandran Suja, Alzheimer’s and Parkinson’s disease therapies in the clinic, Bioeng Transl Med, 2023, 8, e10367.

3. G. E. Vazquez-Velez and H. Y. Zoghbi, Parkinson’s Disease Genetics and Pathophysiology, Annu Rev Neurosci, 2021, 44, 87–108.

4. C. Varadi, Clinical Features of Parkinson’s Disease: The Evolution of Critical Symptoms, Biology (Basel), 2020, 9.

5. B. R. Bloem, M. S. Okun and C. Klein, Parkinson’s disease, Lancet, 2021, 397, 2284–2303.

6. L. V. Kalia and A. E. Lang, Parkinson’s disease, Lancet, 2015, 386, 896–912.

7. A. Recasens and B. Dehay, Alpha-synuclein spreading in Parkinson’s disease, Front Neuroanat, 2014, 8, 159.

8. P. Calabresi, A. Mechelli, G. Natale, L. Volpicelli-Daley, G. Di Lazzaro and V. Ghiglieri, Alpha-synuclein in Parkinson’s disease and other synucleinopathies: from overt neurodegeneration back to early synaptic dysfunction, Cell Death Dis, 2023, 14, 176.

9. P. Calabresi, G. Di Lazzaro, G. Marino, F. Campanelli and V. Ghiglieri, Advances in understanding the function of alpha-synuclein: implications for Parkinson’s disease, Brain, 2023, 146, 3587–3597.

10. L. Stefanis, alpha-Synuclein in Parkinson’s disease, Cold Spring Harb Perspect Med, 2012, 2, a009399.

11. J. Burre, The Synaptic Function of alpha-Synuclein, J Parkinsons Dis, 2015, 5, 699–713.

12. J. Burre, S. Vivona, J. Diao, M. Sharma, A. T. Brunger and T. C. Sudhof, Properties of native brain alpha-synuclein, Nature, 2013, 498, E4–6; discussion E6-7.

13. H. Shen, M. Pirruccello and P. De Camilli, SnapShot: membrane curvature sensors and generators, Cell, 2012, 150, 1300, 1300 e1301–1302.

14. D. H. Johnson and W. F. Zeno, Dynamic interactions between alpha-synuclein and curved membrane surfaces, Biophysical Journal, 2023, 122, 313a.

15. L. Y. Tan, K. H. Tang, L. Y. Y. Lim, J. X. Ong, H. Park and S. Jung, alpha-Synuclein at the Presynaptic Axon Terminal as a Double-Edged Sword, Biomolecules, 2022, 12.

16. M. Vicario, D. Cieri, M. Brini and T. Cali, The Close Encounter Between Alpha-Synuclein and Mitochondria, Front Neurosci, 2018, 12, 388.

17. J. J. Hoozemans, E. S. van Haastert, P. Eikelenboom, R. A. de Vos, J. M. Rozemuller and W. Scheper, Activation of the unfolded protein response in Parkinson’s disease, Biochem Biophys Res Commun, 2007, 354, 707–711.

18. N. Gosavi, H. J. Lee, J. S. Lee, S. Patel and S. J. Lee, Golgi fragmentation occurs in the cells with prefibrillar alpha-synuclein aggregates and precedes the formation of fibrillar inclusion, J Biol Chem, 2002, 277, 48984–48992.

19. N. Thayanidhi, J. R. Helm, D. C. Nycz, M. Bentley, Y. Liang and J. C. Hay, Alpha-synuclein delays endoplasmic reticulum (ER)-to-Golgi transport in mammalian cells by antagonizing ER/Golgi SNAREs, Mol Biol Cell, 2010, 21, 1850–1863.

20. L. Maroteaux, J. T. Campanelli and R. H. Scheller, Synuclein: a neuron-specific protein localized to the nucleus and presynaptic nerve terminal, J Neurosci, 1988, 8, 2804–2815.

21. Y. Singh, P. C. Sharpe, H. N. Hoang, A. J. Lucke, A. W. McDowall, S. P. Bottomley and D. P. Fairlie, Amyloid formation from an alpha-helix peptide bundle is seeded by 3(10)-helix aggregates, Chemistry, 2011, 17, 151–160.

22. C. Galvagnion, A. K. Buell, G. Meisl, T. C. Michaels, M. Vendruscolo, T. P. Knowles and C. M. Dobson, Lipid vesicles trigger alpha-synuclein aggregation by stimulating primary nucleation, Nat Chem Biol, 2015, 11, 229–234.

23. D. Kurouski, Elucidating the Role of Lipids in the Aggregation of Amyloidogenic Proteins, Acc Chem Res, 2023, 56, 2898–2906.

24. B. A. Hijaz and L. A. Volpicelli-Daley, Initiation and propagation of alpha-synuclein aggregation in the nervous system, Mol Neurodegener, 2020, 15, 19.

25. R. Guerrero-Ferreira, N. M. Taylor, D. Mona, P. Ringler, M. E. Lauer, R. Riek, M. Britschgi and H. Stahlberg, Cryo-EM structure of alpha-synuclein fibrils, Elife, 2018, 7.

26. P. Alam, L. Bousset, R. Melki and D. E. Otzen, alpha-synuclein oligomers and fibrils: a spectrum of species, a spectrum of toxicities, J Neurochem, 2019, 150, 522–534.

27. A. Bigi, R. Cascella and C. Cecchi, alpha-Synuclein oligomers and fibrils: partners in crime in synucleinopathies, Neural Regen Res, 2023, 18, 2332–2342.

28. G. M. Compagnoni, A. Di Fonzo, S. Corti, G. P. Comi, N. Bresolin and E. Masliah, The Role of Mitochondria in Neurodegenerative Diseases: the Lesson from Alzheimer’s Disease and Parkinson’s Disease, Mol Neurobiol, 2020, 57, 2959–2980.

29. F. A. Bustamante-Barrientos, N. Luque-Campos, M. J. Araya, E. Lara-Barba, J. de Solminihac, C. Pradenas, L. Molina, Y. Herrera-Luna, Y. Utreras-Mendoza, R. Elizondo-Vega, A. M. Vega-Letter and P. Luz-Crawford, Mitochondrial dysfunction in neurodegenerative disorders: Potential therapeutic application of mitochondrial transfer to central nervous system-residing cells, J Transl Med, 2023, 21.

30. M. T. Henrich, W. H. Oertel, D. J. Surmeier and F. F. Geibl, Mitochondrial dysfunction in Parkinson’s disease - a key disease hallmark with therapeutic potential, Mol Neurodegener, 2023, 18, 83.

31. A. Bose and M. F. Beal, Mitochondrial dysfunction in Parkinson’s disease, J Neurochem, 2016, 139 Suppl 1, 216–231.

32. P. R. Angelova and A. Y. Abramov, Role of mitochondrial ROS in the brain: from physiology to neurodegeneration, FEBS Lett, 2018, 592, 692–702.

33. K. Debnath, A. K. Sarkar, N. R. Jana and N. R. Jana, Inhibiting Protein Aggregation by Small Molecule-Based Colloidal Nanoparticles, Accounts Mater Res, 2022, 3, 54–66.

34. R. Rajan, S. Ahmed, N. Sharma, N. Kumar, A. Debas and K. Matsumura, Review of the current state of protein aggregation inhibition from a materials chemistry perspective: special focus on polymeric materials, Mater Adv, 2021, 2.

35. V. L. Ngoungoure, J. Schluesener, P. F. Moundipa and H. Schluesener, Natural polyphenols binding to amyloid: a broad class of compounds to treat different human amyloid diseases, Mol Nutr Food Res, 2015, 59, 8–20.

36. S. Abrahams, H. C. Miller, C. Lombard, F. H. van der Westhuizen and S. Bardien, Curcumin pre-treatment may protect against mitochondrial damage in LRRK2-mutant Parkinson’s disease and healthy control fibroblasts, Biochem Biophys Rep, 2021, 27, 101035.

37. W. W. Wang, R. Han, H. J. He, J. Li, S. Y. Chen, Y. Gu and C. Xie, Administration of quercetin improves mitochondria quality control and protects the neurons in 6-OHDA-lesioned Parkinson’s disease models, Aging (Albany NY), 2021, 13, 11738–11751.

38. A. Negida, A. Menshawy, G. El Ashal, Y. Elfouly, Y. Hani, Y. Hegazy, S. El Ghonimy, S. Fouda and Y. Rashad, Coenzyme Q10 for Patients with Parkinson’s Disease: A Systematic Review and Meta-Analysis, CNS Neurol Disord Drug Targets, 2016, 15, 45–53.

39. H. Wang, X. Dong, Z. Liu, S. Zhu, H. Liu, W. Fan, Y. Hu, T. Hu, Y. Yu, Y. Li, T. Liu, C. Xie, Q. Gao, G. Li, J. Zhang, Z. Ding and J. Sun, Resveratrol Suppresses Rotenone-induced Neurotoxicity Through Activation of SIRT1/Akt1 Signaling Pathway, Anat Rec (Hoboken), 2018, 301, 1115–1125.

40. I. L. Cockburn, E. R. Pesce, J. M. Pryzborski, M. T. Davies-Coleman, P. G. K. Clark, R.A. Keyzers, L. L. Stephens and G. L. Blatch, Screening for small molecule modulators of Hsp70 chaperone activity using protein aggregation suppression assays: inhibition of the plasmodial chaperone PfHsp70-1, Biol Chem, 2011, 392, 431–438.

41. M. Brys, L. Fanning, S. Hung, A. Ellenbogen, N. Penner, M. Yang, M. Welch, E. Koenig, E. David, T. Fox, S. Makh, J. Aldred, I. Goodman, B. Pepinsky, Y. Liu, D. Graham, A. Weihofen and J. M. Cedarbaum, Randomized phase I clinical trial of anti-alpha-synuclein antibody BIIB054, Mov Disord, 2019, 34, 1154–1163.

42. Jankovic, I. Goodman, B. Safirstein, T. K. Marmon, D. B. Schenk, M. Koller, W. Zago, D. K. Ness, S. G. Griffith, M. Grundman, J. Soto, S. Ostrowitzki, F. G. Boess, M. Martin-Facklam, J. F. Quinn, S. H. Isaacson, O. Omidvar, A. Ellenbogen and G. G. Kinney, Safety and Tolerability of Multiple Ascending Doses of PRX002/RG7935, an Anti-alpha-Synuclein Monoclonal Antibody, in Patients With Parkinson Disease: A Randomized Clinical Trial, JAMA Neurol, 2018, 75, 1206–1214.

43. A. Mahboob, H. Ali, A. AlNaimi, M. Yousef, M. Rob, N. A. Al-Muhannadi, D. K. L. Senevirathne and A. Chaari, Immunotherapy for Parkinson’s Disease and Alzheimer’s Disease: A Promising Disease-Modifying Therapy, Cells, 2024, 13.

44. V. Giorgio, M. Schiavone, C. Galber, M. Carini, T. Da Ros, V. Petronilli, F. Argenton, V. Carelli, M. J. Acosta Lopez, L. Salviati, M. Prato and P. Bernardi, The idebenone metabolite QS10 restores electron transfer in complex I and coenzyme Q defects, Biochim Biophys Acta Bioenerg, 2018, 1859, 901–908.

45. A. Yan, Z. Liu, L. Song, X. Wang, Y. Zhang, N. Wu, J. Lin, Y. Liu and Z. Liu, Idebenone Alleviates Neuroinflammation and Modulates Microglial Polarization in LPS-Stimulated BV2 Cells and MPTP-Induced Parkinson’s Disease Mice, Front Cell Neurosci, 2018, 12, 529.

46. You, M. D. Hossain and Z. Luo, Synthesis of 2D transition metal dichalcogenides by chemical vapor deposition with controlled layer number and morphology, Nano Converg, 2018, 5, 26.

47. Y. Hwang and N. Shin, Colloidal Synthesis of MoSe(2)/WSe(2) Heterostructure Nanoflowers via Two-Step Growth, Materials (Basel), 2021, 14.

48. R. Canton-Vitoria, T. Hotta, M. Xue, S. Zhang and R. Kitaura, Synthesis and Characterization of Transition Metal Dichalcogenide Nanoribbons Based on a Controllable O(2) Etching, JACS Au, 2023, 3, 775–784.

49. H. Ganesha, S. Veeresh, Y. S. Nagaraju, M. Vandana, M. Basappa, H. Vijeth and H. Devendrappa, 2-Dimensional layered molybdenum disulfide nanosheets and CTAB-assisted molybdenum disulfide nanoflower for high performance supercapacitor application, Nanoscale Adv, 2022, 4, 521–531.

50. Y. Jiang, Y. Kang, J. Liu, S. Yin, Z. Huang and L. Shao, Nanomaterials alleviating redox stress in neurological diseases: mechanisms and applications, J Nanobiotechnology, 2022, 20, 265.

51. C. L. Mitchell, M. Matveyenka and D. Kurouski, Neuroprotective properties of transition metal dichalcogenide nanoflowers alleviate acute and chronic neurological conditions linked to mitochondrial dysfunction, J Biol Chem, 2025, 301, 108498.

52. J. Shi, J. Li, Y. Wang, J. Cheng and C. Y. Zhang, Recent advances in MoS(2)-based photothermal therapy for cancer and infectious disease treatment, J Mater Chem B, 2020, 8, 5793–5807.

53. S. K. Ke, P. Y. Yang, Y. G. Wang, S. F. Ye and S. B. Kou, Flower-Like Molybdenum Disulfide Nanostructures for Promoting Mitochondrial Homeostasis and Attenuating Inflammatory Endothelial Dysfunction, Acs Appl Nano Mater, 2021, 4, 11709–11722.

54. X. Qi, L. Li, P. Ye and M. Xie, Macrophage Membrane-Modified MoS(2) Quantum Dots as a Nanodrug for Combined Multi-Targeting of Alzheimer’s Disease, Adv Healthc Mater, 2024, 13, e2303211.

55. G. L. Sutphin and M. Kaeberlein, Measuring Caenorhabditis elegans life span on solid media, J Vis Exp, 2009, DOI: 10.3791/1152.

56. X. Wang, K. Becker, N. Levine, M. Zhang, A. P. Lieberman, D. J. Moore and J. Ma, Pathogenic alpha-synuclein aggregates preferentially bind to mitochondria and affect cellular respiration, Acta Neuropathol Commun, 2019, 7, 41.

57. S. Parihar, A. Parihar, M. Fujita, M. Hashimoto and P. Ghafourifar, Alpha-synuclein overexpression and aggregation exacerbates impairment of mitochondrial functions by augmenting oxidative stress in human neuroblastoma cells, Int J Biochem Cell Biol, 2009, 41, 2015–2024.

58. A. Lurette, R. Martin-Jimenez, M. Khan, R. Sheta, S. Jean, M. Schofield, M. Teixeira, R. Rodriguez-Aller, I. Perron, A. Oueslati and E. Hebert-Chatelain, Aggregation of alpha-synuclein disrupts mitochondrial metabolism and induce mitophagy via cardiolipin externalization, Cell Death Dis, 2023, 14, 729.

59. Shen and U. Dettmer, Alpha-Synuclein Effects on Mitochondrial Quality Control in Parkinson’s Disease, Biomolecules, 2024, 14.

60. J. H. Park, J. D. Burgess, A. H. Faroqi, N. N. DeMeo, F. C. Fiesel, W. Springer, M. Delenclos and P. J. McLean, Alpha-synuclein-induced mitochondrial dysfunction is mediated via a sirtuin 3-dependent pathway, Mol Neurodegener, 2020, 15, 5.

61. B. J. Ryan, S. Hoek, E. A. Fon and R. Wade-Martins, Mitochondrial dysfunction and mitophagy in Parkinson’s: from familial to sporadic disease, Trends Biochem Sci, 2015, 40, 200–210.

62. A. Wilkaniec, A. M. Lenkiewicz, L. Babiec, E. Murawska, H. M. Jesko, M. Cieslik, C. Culmsee and A. Adamczyk, Exogenous Alpha-Synuclein Evoked Parkin Downregulation Promotes Mitochondrial Dysfunction in Neuronal Cells. Implications for Parkinson’s Disease Pathology, Front Aging Neurosci, 2021, 13, 591475.

63. D. Freeman, R. Cedillos, S. Choyke, Z. Lukic, K. McGuire, S. Marvin, A. M. Burrage, S. Sudholt, A. Rana, C. O’Connor, C. M. Wiethoff and E. M. Campbell, Alpha-synuclein induces lysosomal rupture and cathepsin dependent reactive oxygen species following endocytosis, PLoS One, 2013, 8, e62143.

64. A. K. Reeve, M. H. Ludtmann, P. R. Angelova, E. M. Simcox, M. H. Horrocks, D. Klenerman, S. Gandhi, D. M. Turnbull and A. Y. Abramov, Aggregated alpha-synuclein and complex I deficiency: exploration of their relationship in differentiated neurons, Cell Death Dis, 2015, 6, e1820.

65. A. Wolozin and P. Ivanov, Stress granules and neurodegeneration, Nat Rev Neurosci, 2019, 20, 649–666.

66. Y. Liu, Z. Yao, G. Lian and P. Yang, Biomolecular phase separation in stress granule assembly and virus infection, Acta Biochim Biophys Sin (Shanghai), 2023, 55, 1099–1118.

67. Q. Cui, Z. Liu and G. Bai, Friend or foe: The role of stress granule in neurodegenerative disease, Neuron, 2024, 112, 2464–2485.

68. S. M. Heman-Ackah, R. Manzano, J. J. M. Hoozemans, W. Scheper, R. Flynn, W. Haerty, S. A. Cowley, A. R. Bassett and M. J. A. Wood, Alpha-synuclein induces the unfolded protein response in Parkinson’s disease SNCA triplication iPSC-derived neurons, Hum Mol Genet, 2017, 26, 4441–4450.

69. A. Bellucci, L. Navarria, M. Zaltieri, E. Falarti, S. Bodei, S. Sigala, L. Battistin, M. Spillantini, C. Missale and P. Spano, Induction of the unfolded protein response by alpha-synuclein in experimental models of Parkinson’s disease, J Neurochem, 2011, 116, 588–605.

70. E. Colla, Linking the Endoplasmic Reticulum to Parkinson’s Disease and Alpha-Synucleinopathy, Front Neurosci, 2019, 13, 560.

71. Siwecka, K. Saramowicz, G. Galita, W. Rozpedek-Kaminska and I. Majsterek, Inhibition of Protein Aggregation and Endoplasmic Reticulum Stress as a Targeted Therapy for alpha-Synucleinopathy, Pharmaceutics, 2023, 15.

72. M. Huang, B. Wang, X. Li, C. Fu, C. Wang and X. Kang, alpha-Synuclein: A Multifunctional Player in Exocytosis, Endocytosis, and Vesicle Recycling, Front Neurosci, 2019, 13, 28.

73. S. Hui, J. George, M. Kapadia, H. Chau, Z. Bariring, R. Earnshaw, K. Shafiq, L. V. Kalia and S. K. Kalia, Mitophagy Upregulation Occurs Early in the Neurodegenerative Process Mediated by alpha-Synuclein, Mol Neurobiol, 2024, 61, 9032–9042.

74. S. Sofela, S. Sahloul and Y. A. Song, Biophysical analysis of drug efficacy on C. elegans models for neurodegenerative and neuromuscular diseases, PLoS One, 2021, 16, e0246496.

75. G. Hernando and C. Bouzat, Drug combination assays using Caenorhabditis elegans as a model system, J Pharmacol Toxicol Methods, 2025, 131, 107583.

76. L. Tejeda-Benitez and J. Olivero-Verbel, Caenorhabditis elegans, a Biological Model for Research in Toxicology, Rev Environ Contam Toxicol, 2016, 237, 1–35.

77. A. Roussos, K. Kitopoulou, F. Borbolis and K. Palikaras, Caenorhabditis elegans as a Model System to Study Human Neurodegenerative Disorders, Biomolecules, 2023, 13.

78. E. Caldero-Escudero, S. Romero-Sanz and S. De la Fuente, Using C. elegans as a model for neurodegenerative diseases: Methodology and evaluation, Methods Cell Biol, 2024, 188, 1–34.

79. A. K. Torres, R. G. Mira, C. Pinto and N. C. Inestrosa, Studying the mechanisms of neurodegeneration: C. elegans advantages and opportunities, Front Cell Neurosci, 2025, 19, 1559151.

80. K. Strange, From genes to integrative physiology: ion channel and transporter biology in Caenorhabditis elegans, Physiol Rev, 2003, 83, 377–415.

81. L. A. Herndon, C. A. Wolkow, M. Driscoll and D. H. Hall, Effects of Ageing on the Basic Biology and Anatomy of. Healthy Ageing Long, 2017, 5, 9–39.

82. A. Jan, N. P. Goncalves, C. B. Vaegter, P. H. Jensen and N. Ferreira, The Prion-Like Spreading of Alpha-Synuclein in Parkinson’s Disease: Update on Models and Hypotheses, Int J Mol Sci, 2021, 22.

83. W. P. Ruf, J. L. Meirelles and K. M. Danzer, Spreading of alpha-synuclein between different cell types, Behav Brain Res, 2023, 436, 114059.

84. J. Thorne and D. A. Tumbarello, The relationship of alpha-synuclein to mitochondrial dynamics and quality control, Front Mol Neurosci, 2022, 15, 947191.

85. F. F. Geibl, M. T. Henrich, Z. Xie, E. Zampese, J. Ueda, T. Tkatch, D. L. Wokosin, E. Nasiri, C. A. Grotmann, V. L. Dawson, T. M. Dawson, N. S. Chandel, W. H. Oertel and D. J. Surmeier, alpha-Synuclein pathology disrupts mitochondrial function in dopaminergic and cholinergic neurons at-risk in Parkinson’s disease, Mol Neurodegener, 2024, 19, 69.

86. H. Zeng, Y. Liu, X. Liu, J. Li, L. Lu, C. Xue, X. Wu, X. Zhang, Z. Zheng and G. Lu, Interplay of alpha-Synuclein Oligomers and Endoplasmic Reticulum Stress in Parkinson’S Disease: Insights into Cellular Dysfunctions, Inflammation, 2025, 48, 1590–1606.

87. H. Xiang, The interplay between alpha-synuclein aggregation and necroptosis in Parkinson’s disease: a spatiotemporal perspective, Front Neurosci, 2025, 19, 1567445.

88. M. R. Glineburg, E. Yildirim, N. Gomez, G. Rodriguez, J. Pak, X. Li, C. Altheim, J. Waksmacki, G. M. McInerney, S. J. Barmada and P. K. Todd, Stress granule formation helps to mitigate neurodegeneration, Nucleic Acids Res, 2024, 52, 9745–9759.

89. A. R. Winslow and D. C. Rubinsztein, The Parkinson disease protein alpha-synuclein inhibits autophagy, Autophagy, 2011, 7, 429–431.

90. Y. B. Mingo, M. L. Escobar Galvis and M. X. Henderson, alpha-Synuclein pathology and mitochondrial dysfunction: Toxic partners in Parkinson’s disease, Neurobiol Dis, 2025, 209, 106889.

91. S. Rius-Perez, I. Torres-Cuevas, I. Millan, A. L. Ortega and S. Perez, PGC-1alpha, Inflammation, and Oxidative Stress: An Integrative View in Metabolism, Oxid Med Cell Longev, 2020, 2020, 1452696.

