## Supplemental Figure 1 for "Neuroprotective Parkinson’s Disease Therapeutic: Transition Metal Dichalcogenide Nanoflower Treatments Alleviate Pathological Cell Stress"

Charles L. Mitchell[1,2], Mikhail Matveyenka[2], Harris C. Brown[2], Jessica Aldape[2], Payton Moore[2], Kha-Tran Nguyen[2], John C. Walker[2], Bryce Pearson[2], Joshua Skrehot[2], Dmitry Kurouski[1,2*]
[1] Interdisciplinary Program in Genetics and Genomics, Texas A&M University, College Station, Texas, USA
[2] Department of Biochemistry and Biophysics, Texas A&M University, College Station, Texas, USA

### Supporting Information

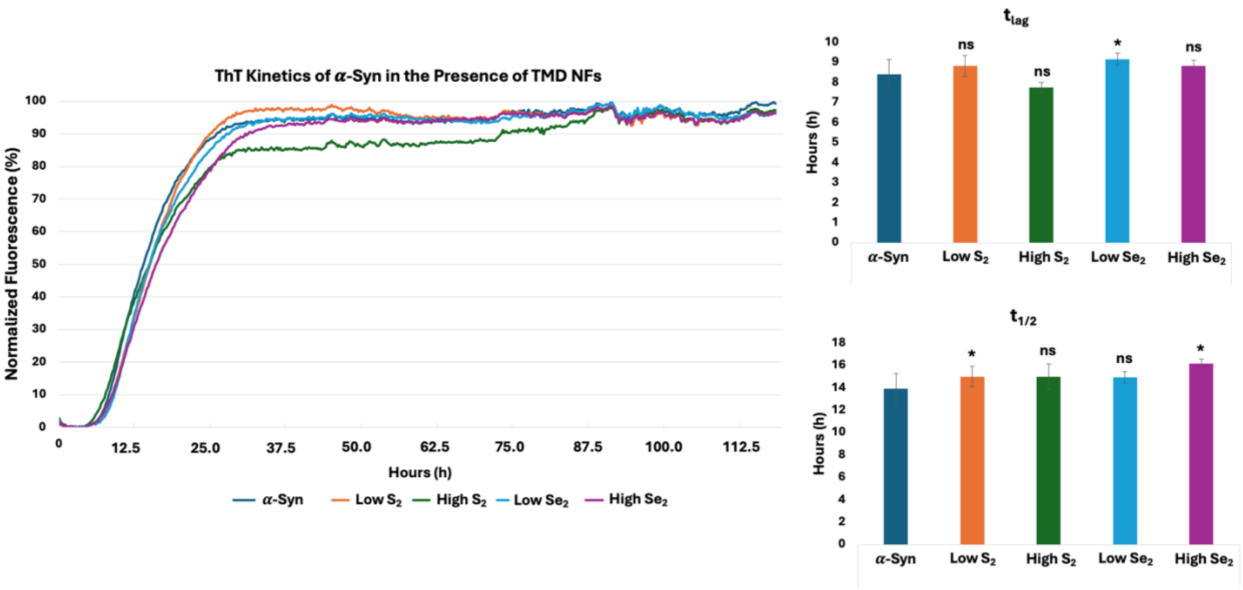


**Figure S1:** Aggregation kinetics for α-syn protein in the presence of TMD NFs at 0.1 mg/mL (low) and 1.0 mg/mL (high) concentrations via fluorescence of thioflavin T (ThT) [left]. Calculated $t_{lag}$ [top right] and $t_{1/2}$ [bottom right] from the aggregation curves for α-syn in the presence of 0.1 mg/mL $MoS_2$ (Low $S_2$), 1.0 mg/mL $MoS_2$ (High $S_2$), 0.1 mg/mL $MoSe_2$ (Low $Se_2$), 1.0 mg/mL $MoSe_2$ (High $Se_2$), and the absence of NFs (α-Syn). Significance marking coincide with comparisons made to the α-syn control.
